# A very-low carbohydrate content in a high-fat diet modifies the plasma metabolome and alleviates experimental atherosclerosis

**DOI:** 10.1101/2023.02.07.527515

**Authors:** R Castro, K Kalecký, NK Huang, K Petersen, V Singh, AC Ross, T Neuberger, T Bottiglieri

## Abstract

Ketogenic diets (KD) are very low-carbohydrate diets that promote nutritional ketosis and are widely used for weight loss, although concerns about potential adverse cardiovascular effects remain. In this study, we used apolipoprotein E deficient (*ApoE* ^*-/-*^) mice to investigate the vascular impact and plasma metabolic signature of a very high-fat KD compared to a non-ketogenic high-fat diet (HFD). Plasma samples were collected after 4, 8, and 12 weeks on the experimental diets and used to quantify the major ketone body β-hydroxybutyrate (BHB), inflammatory cytokines (interleukin 6, IL-6; monocyte chemoattractant protein 1, MCP-1; and tumor necrosis factor α, TNF-α), and targeted metabolomic profiling by mass spectrometry. Moreover, aortic atherosclerotic lesions were quantified ex vivo by magnetic resonance imaging (MRI) on a 14-tesla system. The results showed that, relative to HFD mice, the KD mice had markedly higher levels of BHB and lower levels of cytokines, confirming the presence of ketosis that alleviated the well-established fat-induced systemic inflammation. Moreover, mice under nutritional ketosis displayed a distinct plasma amino acid profile evidencing a KD-induced alteration in protein metabolism. Significant changes in the plasma metabolome in KD mice included a decrease in lipophilic and increase in hydrophilic metabolites. Despite the higher fat content of the KD versus the HFD, KD mice presented significantly lower levels of several lipid metabolites, including phosphatidylcholines, cholesterol esters, sphingomyelins, and ceramides. Consistent with the shift in energy metabolism toward fatty acid oxidation caused by the KD, the ratio of acylcarnitines to free carnitine was significantly higher in KD than in HFD mice., the aortic plaque burden was significantly lower in the KD versus the HFD group. In conclusion, nutritional ketosis induced by the KD was associated with specific metabolic changes and an atheroprotective phenotype versus the HFD.

## Introduction

Cardiovascular diseases (CVDs) are the leading cause of death globally; about 659,000 people in the United States die from CVD each year ^1^. The major underlying cause of most CVD is atherosclerosis, a multifactorial chronic inflammatory disease of the arteries ^2^. Atherosclerosis is characterized by the accumulation of lipids, inflammatory factors, and plaque in the arterial walls. Nutrition has a central role in both primary and secondary prevention of CVD ^3-5^. Adherence to a heart-healthy dietary pattern is associated with lower risk of CVD^5^. Heart-healthy dietary patterns emphasize fruits, vegetables, wholegrains, nuts, seeds, legumes, and lean protein sources. These dietary patterns typically have a macronutrient distribution consistent with the Acceptable Macronutrient Distribution Range (AMDR; carbohydrates 45-65% kcal; fat 20-35% kcal; protein 10-35% kcal)^6^. There is currently interest in identifying potential health benefits, including cardioprotection, of dietary patterns with a macronutrient composition outside the AMDR^7^.

Ketogenic diets (KD) are very-high fat diets (>70% kcal) that are being increasingly used for weight loss; nevertheless, concerns about possible fat-induced adverse cardiovascular effects exist ^8,9^. Severe restriction of carbohydrate consumption, with a concurrent increase in fat, results in the promotion of lipid oxidation to produce ketone bodies, and thus, a metabolic state of nutritional ketosis ^10^. The ketone body β-hydroxybutyrate (BHB) is synthesized in the liver from fatty acids and represents an essential carrier of energy from the liver to peripheral tissues when the supply of glucose is too low for the body’s energetic needs. Interestingly, in addition to its activity as an energetic metabolite, BHB is increasingly understood to have signaling functions that may be relevant to a variety of human diseases including CVD ^11^.

The composition of KD that are capable of producing nutritional ketosis is typically outside the range of diets considered cardioprotective. To achieve nutritional ketosis, KDs are typically inconsistent with nutritional recommendations for CVD prevention raising vascular safety concerns; in fact, cardio-protective foods are severely restricted (e.g., fruits, legumes) whereas foods associated with increased CVD risk are promoted (e.g., meats rich in saturated fat) ^8,9,12^

In contrast to this notion that higher fat in KD might heighten the levels of CVD risk factors, both preclinical and experimental human studies demonstrated the beneficial effects of KD in improving multiple components of metabolic syndrome, an established risk factor of CVD. More importantly, a recent study documents that KD extends longevity and health span in mice ^13^. Therefore, to test whether KD exacerbate or attenuate atherosclerosis development, we employed the apolipoprotein E deficient (*ApoE*−/ −) mouse model, which is a well characterized pre-clinical model to study all recognized stages of human atherosclerosis ^14^. In addition, the complexity and morphological features of atherosclerotic lesions that develop in this mouse model are very similar to those in humans ^14^. Atherosclerosis in *ApoE*−/ − mice is driven by impaired clearance of cholesterol-enriched lipoproteins, which results in elevated levels of plasma cholesterol and atherogenic remnants.

In the present study the vascular effect of a very high-fat very-low carbohydrate KD compared to a non-ketogenic high-fat diet with higher carbohydrate content was determined as well as the corresponding systemic metabolic signatures. With that purpose, *ApoE*^−/ −^ mice were fed two high-fat diets (Kcal%, carbohydrate/fat/protein), specifically, KD (1/81/18,) or HFD (42/40/18), or a control diet in order to investigate atherogenic, inflammatory markers and metabolic changes.

## 2. Materials and Methods

### 2.1. Animals and Diets

Seven-week-old *ApoE*^−/ −^ mice, purchased from Jackson Laboratory (Bar Harbor, ME, USA) were housed in a temperature- and humidity-controlled room. Only male mice were used to control for the known effect of gender on atherosclerosis in this strain ^15^. The animals were housed individually and had free access to water and to one of the following diets prepared based on AIN 93G (Research Diets, New Brunswick, NJ, USA): a control diet (11 Kcal% fat, 70 Kcal% carbohydrate, 18 Kcal% protein), a high-fat diet (HFD) (41 Kcal% fat, 40 Kcal% carbohydrate, 18 Kcal% protein and 0.15% cholesterol) or a KD diet (81 Kcal% fat, 1 Kcal% carbohydrate, 18 Kcal% protein and 0.15% cholesterol). All three diets contained the same levels of micronutrients. Diets were replaced once a week, at which time animals and the remaining food were weighed to determine food consumption and body weight progression. All procedures were performed in compliance with the Institutional Animal Care and Use Committee of the Pennsylvania State University, which approved this study.

### 2.2. Blood Collection

After mice were fed for 4, 8 and 12 weeks, blood was collected from the retro-orbital cavity into heparinized tubes and immediately placed on ice. Plasma was isolated by centrifugation at 4°C and immediately stored at −80°C prior to further biochemical and metabolomic analyses.

### 2.3. Triacylglycerols, Total Cholesterol, β-Hydroxybutyrate, and Glucose

Plasma collected at 8 weeks was tested for triacylglycerols (TAG), and total cholesterol contents using colorimetric assays kits (Randox, Ann Arbor, MI, USA) following the manufacturer’s instructions.

Plasma collected at 4, 8, and 12 weeks was tested for the concentration of BHB and glucose. BHB was determined using a colorimetric assay kit (Cayman Chemical, Ann Arbor, MI, USA) following the manufacturer’s instructions. Blood glucose was measured using a glucometer (Contour, Bayer, Tarrytown, NY, USA) following the manufacturer’s instructions.

### 2.4. Metabolomic Analysis

Targeted metabolomics in plasma collected at 8 weeks was performed using the commercially available MxP Quant 500 kit (Biocrates life sciences, Innsbruck, Austria), following the manufacturer’s instructions provided. 10 μL of plasma, calibrators (7 levels) and quality controls (3 levels) were added to the 96 well kit plates. The MxP® Quant 500 kit can potentially identify 630 metabolites across 23 classes of compounds by liquid chromatography (LC) and flow-injection analysis (FIA) coupled to tandem mass spectrometry (MS/MS), and calculate additional 232 metabolic indicators based on sums and ratios of metabolites.

### 2.5. Proinflammatory cytokines

Plasma collected at 12 weeks, was used to determine the concentration of the proinflammatory cytokines, IL-6 Interleukin 6 (IL-6), monocyte chemoattractant protein 1 (MCP-1), and tumor necrosis factor α (TNF-α) by ELISA (Meso Scale Diagnostics, Rockville, MD, USA) following the manufacturer’s instructions.

### 2.6. Aorta Collection and 14T-MRI Analysis

After 12 weeks, mice were euthanized by carbon dioxide inhalation and aortas were collected, as previously described in detail ^16-18^. Briefly, aortas were perfused with 10 mL of cold phosphate saline buffer (PBS), followed by 10% neutral buffered formalin (NBF) in PBS, and were then fixed overnight in 10% NBF. The aortas were further equilibrated in a solution of 0.1% Magnevist (Bayer, Whippany, NJ, USA), 0.25% sodium azide in PBS overnight at 4°C, and next transferred into glass tubes (6 mm OD, 4 mm ID and 60 mm length) for magnetic resonance imaging (MRI) analysis. Aortic plaque burden was then quantified using an Agilent 14T micro imaging system, as previously described in detail ^16-18^. A gradient echo imaging sequence with an imaging time of 9 h 48 min was used to generate 3D datasets of the aortas. Scan parameters included an echo time (TE) of 13 ms, a repetition time (TR) of 100.00 ms, eight averages, a field of view (FOV) of 12.6 × 4.2 × 4.2 mm^3^, and a matrix size of 630 × 210 × 210, resulting in an isotropic resolution of 20 μm. After acquisition, MR data were reconstructed using MATLAB (The MathWorks Inc., Natick, MA, USA). Zero-filling by a factor of 2 in each direction led to a final isotropic pixel resolution of 10 μm.

Data segmentation was performed using Avizo 9.5 (Thermo Fisher Scientific, Waltham, MA, USA). The lumen of the aorta, the different plaques and the aorta wall were manually segmented using the lasso tool by two blinded subjects. Quantification of plaque volume was determined using the material statistics function in Avizo 9.5 on the segmented aorta. Different segments of the vessel were considered, namely, whole aorta, aortic arch and brachiocephalic artery. Results were expressed as the percent of plaque area in relation to the total segmented area (plaque, lumen, and wall).

### 2.7. Statistical analysis and bioinformatics

All analyses, except plasma metabolomics, were performed in GraphPad Prism 7 (GraphPad Software, La Jolla, CA, USA), with statistical significance set to p< 0.05. For comparison of two groups, an unpaired Student’s t-test was used. For more than two groups, a one- or two-way analysis of variance (ANOVA) was performed and adjusted for multiple comparisons.

Plasma metabolomics processing: peak detection, concentration computation, and calculation of metabolic indicators were done in MetIDQ Oxygen v3005 (Biocrates Life Sciences, Innsbruck, Austria). Limits of detection (LODs) were calculated as 3x median signal in phosphate-buffered saline blanks in each plate. Analytes (metabolites and indicators) with more than 50% of samples below LOD in each group were filtered out. Quality control samples were used for median plate normalization. Values were log transformed for analytes only where the transformation improved normality of the distribution as judged by the Shapiro-Wilk test for normality ^19^. Outliers were adjusted via Tukey’s fencing ^20^.

Metabolomics data were analyzed in R v3.6.1 ^21^ and RStudio v1.2.5033 ^22^ through a series of linear regression model with analyte values as regressands and compared groups as regressors with plate information as an additional covariate. Multiple testing was controlled with false discovery rate (FDR) using Benjamini-Hochberg procedure ^23^ and results with FDR ≤ 0.05 were considered significant. Heatmaps were rendered with R package *gplots* v3.0.3 ^24^, using *ward*.*D2* method for clustering. BoxPlots were created with R package *ggplot2* v3.3.1 ^25^. Chemical cluster analysis was performed with online tool ChemRICH ^26^.

## 3. Results

### KD protected from fat-induced weight gain

KD are energy dense diet and animals require relatively low amount of food to meet their energy requirement. In line, we did observe that KD-fed mice consumed significantly less food than mice fed the control or HF diets (Figure 1A). Nevertheless, due to the higher energy density of the KD (6.2 kcal/g KD versus 4.6 kcal/g HFD versus 3.9 kcal/g control diet), these mice consumed more calories than controls and the same number as the HF mice (Figure 1B).

**Figure 1.**
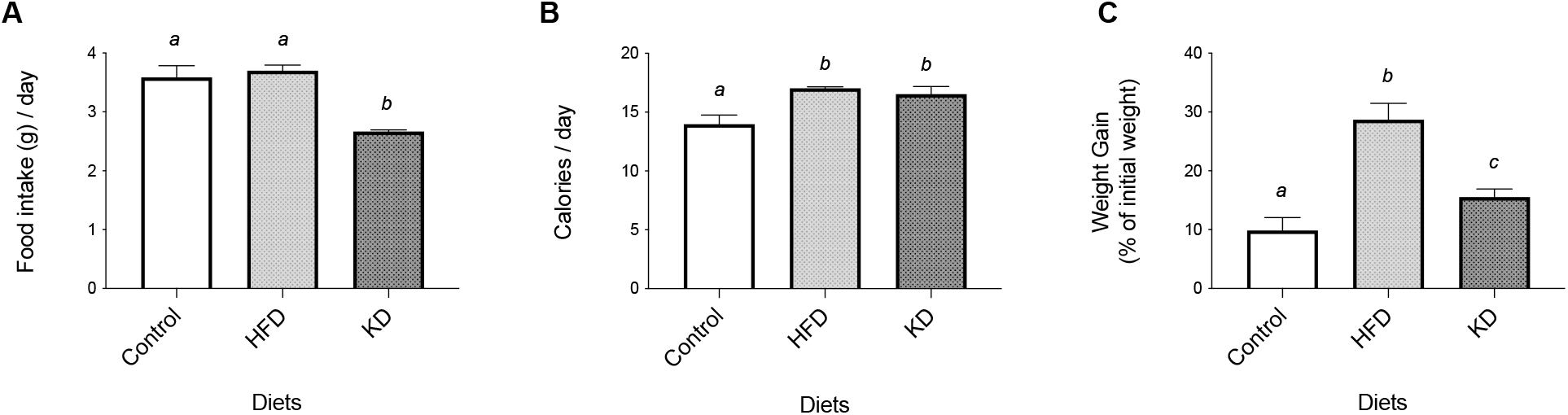
Effect of the experimental diets (control, high-fat diet, HFD, and ketogenic diet, KD) on the daily intake of food (A) and calories (B), and on body weight gain (C). Values are mean ± SD; n= 8-9/group, bars not sharing superscript letters differ from each other, p < 0.05.

Compared to the control diet, as expected, the HF-fed group had significant higher body weight. Notably, however, despite the similar number of calories consumed by both HF- and KD-groups, mice from the KD group weighed significantly less than HF group (Figure 1C).

### KD induced nutritional ketosis, and diminished plasma glucose

The plasma levels of BHB and glucose were measured at three different time points of the feeding trial (after 4, 8, and 12 weeks). At each timepoints tested in this study, BHB concentration was always significantly higher in KD mice than control diet or HF diet-fed groups, confirming the presence of a sustained nutritional ketosis under this dietary condition (Figure 2A). Plasma glucose showed opposite results, with KD-fed mice presenting significantly lower concentrations than controls or HFD-fed mice (Figure 2B). On the other hand, plasma TAG (Figure 2C) and total cholesterol levels (Figure 2D) after 8 weeks on the experimental diets were higher in the HFD-fed mice and lower in controls, with the KD group showing intermediate values for both parameters (Figure 2D).

**Figure 2.**
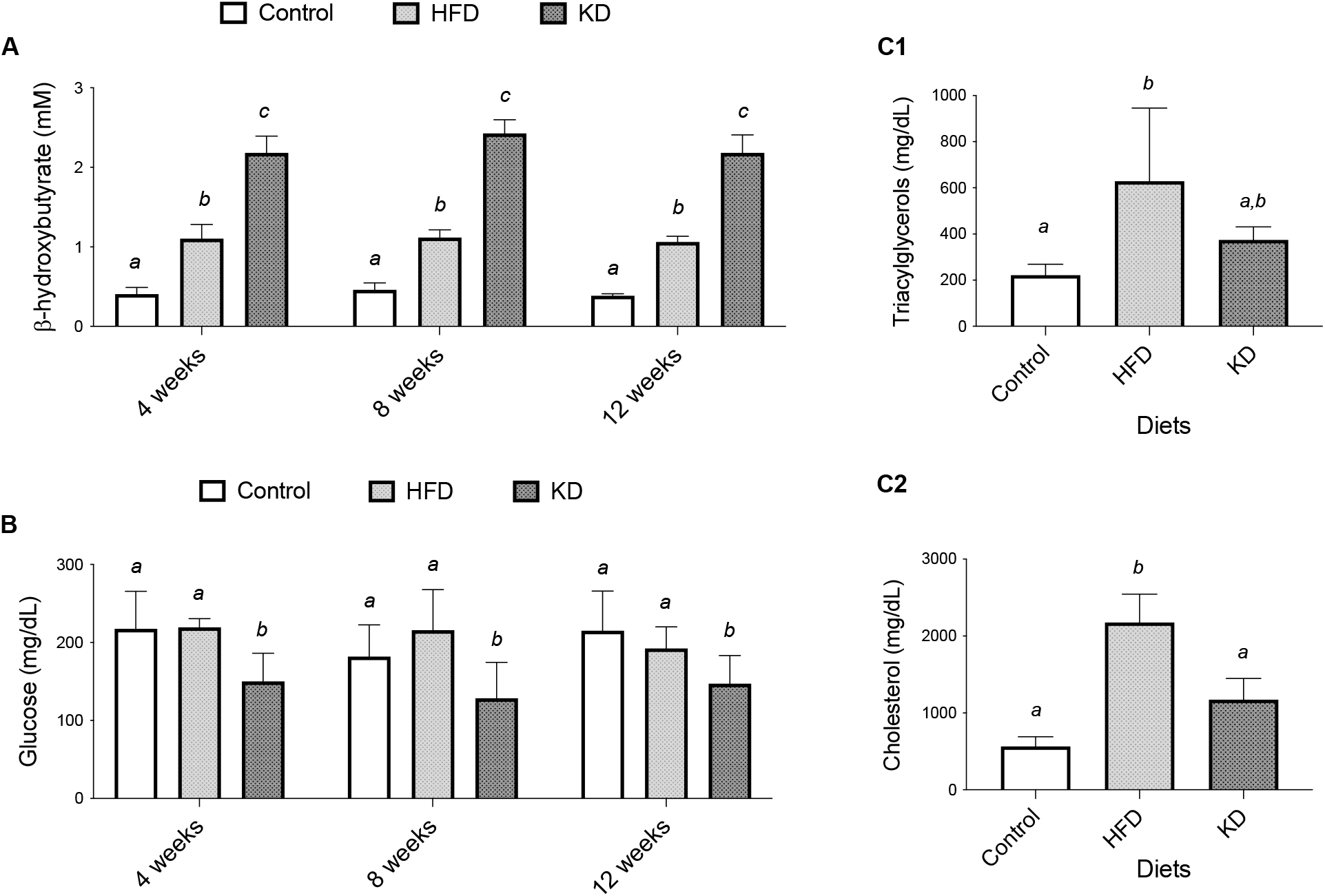
Effect of the experimental diets (control, high-fat diet, HFD, and ketogenic diet, KD) on the circulating levels of the ketone body, ß-hydroxybutyrate (A) and glucose (B) after four, eight, or twelve weeks, and on the levels of triacylglycerols (C1) and total cholesterol (C2) after eight weeks. Values are mean ± SD; n= 6-8/group, bars not sharing superscript letters differ from each other, p < 0.05.

### KD alleviated inflammation and delayed the progression of atherosclerosis

The effect of 12 weeks of the experimental diets on the status of systemic inflammation, a key component of the atherosclerosis process, was also evaluated. The plasma concentrations of the three pro-inflammatory, MCP1 (Figure 3A top), IL-6 (Figure 3A middle), and TNFα (Figure 3A bottom) were determined. The results showed that HF mice presented the highest concentrations of the three cytokines whereas the lowest values were observed in the control-group. This observation agrees with the positive effect of dietary fat on systemic inflammation ^27^. Notably, despite the higher fat content in the KD (80%), than the HFD (40%), KD-fed mice exhibited significantly lower levels of systemic inflammation than the HFD group evidencing a protective role of nutritional ketosis in fat-induced inflammation.

**Figure 3.**
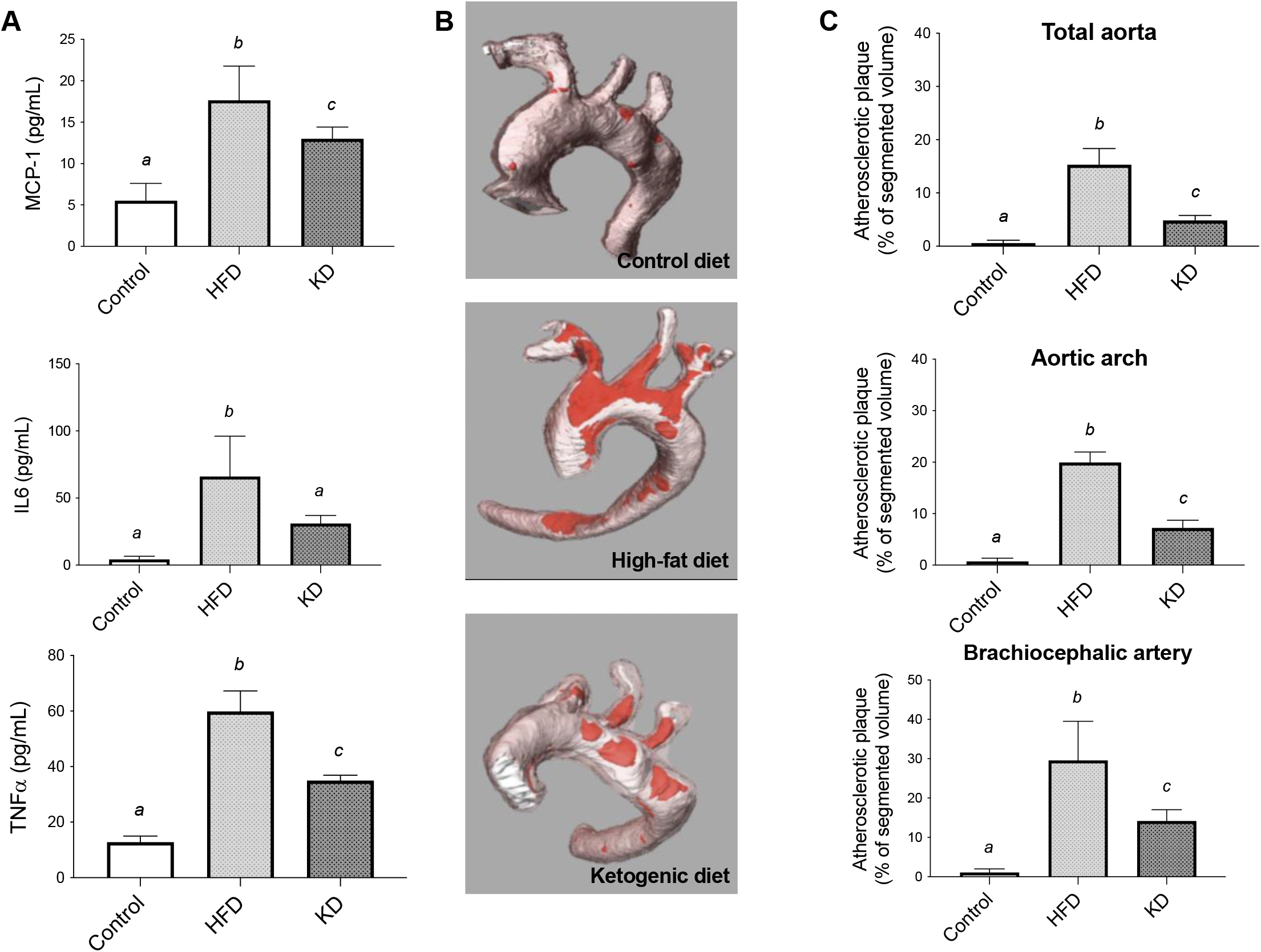
Effect of 12 weeks of the experimental diets (control, high-fat diet, HFD, and ketogenic diet, KD) on systemic inflammation and aortic atherosclerosis assessed by 14T-MRI. A. Levels of monocyte chemoattractant protein 1 (MCP1) (B1), interleukin 6 (IL-6) (B2), and tumor necrosis factor α (TNF-α); B. Representative images of plaque burden from each dietary group (plaque is colored in red); C. % of plaque volume in different aortic segments. Values are mean ± SD; n= 6-8/group, bars not sharing superscript letters differ from each other, p < 0.05.

In this study, high-field magnetic resonance imaging (14T-MRI) was used to quantify the vascular atheroma in aortas obtained from *ApoE*^−/ −^ mice. The results showed that mice fed the control diet presented significantly (*p < 0*.*05*) less aortic atherosclerosis than KD or HF mice, indicating the well-established role of dietary fat in the development of atherosclerosis.

However, significant differences were detected between the latter two groups at the *p < 0*.*05* level (Figure 3). As referred to above, the fat content in both diets was profoundly different (KD, 80%; HFD, 40%). However, in accordance with the lower systemic inflammation detected, the volume of vascular lesions quantified in the aortas of KD-fed mice was significantly (p<0.05) lower than in the HFD group, suggesting an anti-atherogenic effect of the KD. The atherosclerotic lesions were concentrated in the aortic arch and brachiocephalic artery in all three groups; thus, the differences in plaque burden between the HFD, KD, and control aortas in these more atheroprone regions, the aortic arch and brachiocephalic artery, were even more pronounced than when considering the entire aorta (Figure 3B).

### KD mice displayed a distinct metabolic profile

Metabolomic profiling offers a novel approach to identify disease biomarkers, as the levels of specific metabolites are a functional readout and critical indicator of human health and disease. Therefore, a targeted metabolomic approach was applied to investigate further the metabolic changes associated with the vascular effects of the experimental diets. After filtering for compounds below detection, we detected 434 metabolites and lipid species out of a potential of 631 that could be potentially determined using the Quant 500 targeted metabolomic platform. Further analysis revealed 123 metabolites and lipid species were significantly different between the KD and HFD groups (Table 2, Supplementary Table S1), thus showing that the absence of dietary carbohydrates profoundly modified the plasma metabolome of the animals. Significantly lower levels of several complex lipid metabolites were observed in the KD group versus HFD (Table 2; Figure 4A). The lipid classes most affected were glycerophosphopids, sphingolipids, cholesterol esters, ceramides and glyceroceramides (mono, di, and tri-hexosylceramides). Some species of triacylglycerols were also decreased. The decrease in cholesterol esters and triacylglycerols in KD versus HFD determined by mass spectrometry is consistent with the analysis for these lipids performed using the colorimetric assays. Interestingly, however, the plasma concentration of small molecules showed the opposite trend. We observed significantly higher levels of 16 amino acids or related metabolites and 4 bile acids in the KD than in the HFD mice (Table 2; Figure 4B, Supplementary Table S1). However, KD-mice presented lower levels of the small molecule p-cresol sulfate than HF-mice. P-Cresol sulfate is a gut-related metabolite that negatively affects cardiometabolic parameters in mice (triggers insulin resistance and redistribution of lipids) ^28^. Figure 5 shows the effect size for the increase and decrease in specific metabolites and lipid species and illustrates the differences in metabolism between KD and HFD relative to the CD. Overall, KD results in an attenuation of various atherogenic lipid species.

**Table 1.** Macronutrient composition of the experimental diets (control; high-fat diet, HFD; and ketogenic diet, KD)

| Macronutrient by gm<br>(400 Kcal total) | Control | HFD | KD |
| --- | --- | --- | --- |
| Casein | 182 | 181 | 181 |
| Corn Starch | 435 | 215 | 0 |
| Maltodextrin 10 | 156 | 92 | 0 |
| Sucrose | 100 | 100 | 0 |
| Cellulose | 35 | 35 | 35 |
| Cocoa Butter | 0 | 153 | 100 |
| Primex (Non Trans-Fat) | 25 | 0 | 232 |
| Corn Oil | 25 | 25 | 25 |
| Cholesterol | 0 | 11 | 11 |
| L-Cystine | 3.0 | 3.0 | 3.0 |

**Table 2.** Number of metabolites and complex lipids significantly different (FDR <0.05) between the ketogenic diet (KD) and high-fat diet (HFD).

| Analyte Class | Quant 500 |  | KD vs. HFD<br>(FDR <0.05 significant) |  |  |
| --- | --- | --- | --- | --- | --- |
|  | Potential | Detected | Total | Increased | Decreased |
| **Metabolites** |  |  |  |  |  |
| Alkaloids | 1 | 0 | 0 | 0 | 0 |
| Amine Oxides | 1 | 1 | 0 | 0 | 0 |
| Amino Acids | 20 | 20 | 8 | 8 | 0 |
| Amino Acids Related | 31 | 23 | 8 | 8 | 0 |
| Bile Acids | 14 | 9 | 4 | 4 | 0 |
| Biogenic Amines | 9 | 5 | 0 | 0 | 0 |
| Carboxylic Acids | 7 | 2 | 0 | 0 | 0 |
| Cresols | 1 | 1 | 1 | 0 | 1 |
| Fatty Acids | 12 | 7 | 2 | 2 | 0 |
| Hormones | 4 | 0 | 0 | 0 | 0 |
| Indoles and Derivatives | 4 | 2 | 0 | 0 | 0 |
| Nucleobases Related | 2 | 0 | 0 | 0 | 0 |
| Sugars | 1 | 1 | 1 | 0 | 1 |
| Vitamins and Cofactors | 1 | 1 | 0 | 0 | 0 |
| **Complex Lipids** |  |  |  |  |  |
| Acylcarnitines | 40 | 12 | 0 | 0 | 0 |
| Glycerophospholipids<br>(Lysophosphatidylcholines &<br>Phosphatidylcholines) | 90 | 89 | 52 | 1 | 51 |
| Sphingolipids | 15 | 15 | 9 | 0 | 9 |
| Cholesterol Esters | 22 | 17 | 8 | 0 | 8 |
| Ceramides | 28 | 18 | 5 | 0 | 5 |
| Dihydroceramides | 8 | 4 | 0 | 0 | 0 |
| Glyceroceramides<br>(Mono-, Di-, and Tri-hexosylceramides) | 34 | 30 | 16 | 0 | 16 |
| Diacylglycerols | 44 | 5 | 1 | 0 | 1 |
| Triacylglycerols | 242 | 172 | 8 | 2 | 6 |
| **Total** | **631** | **434** | **123** | **25** | **98** |

**Figure 4.**
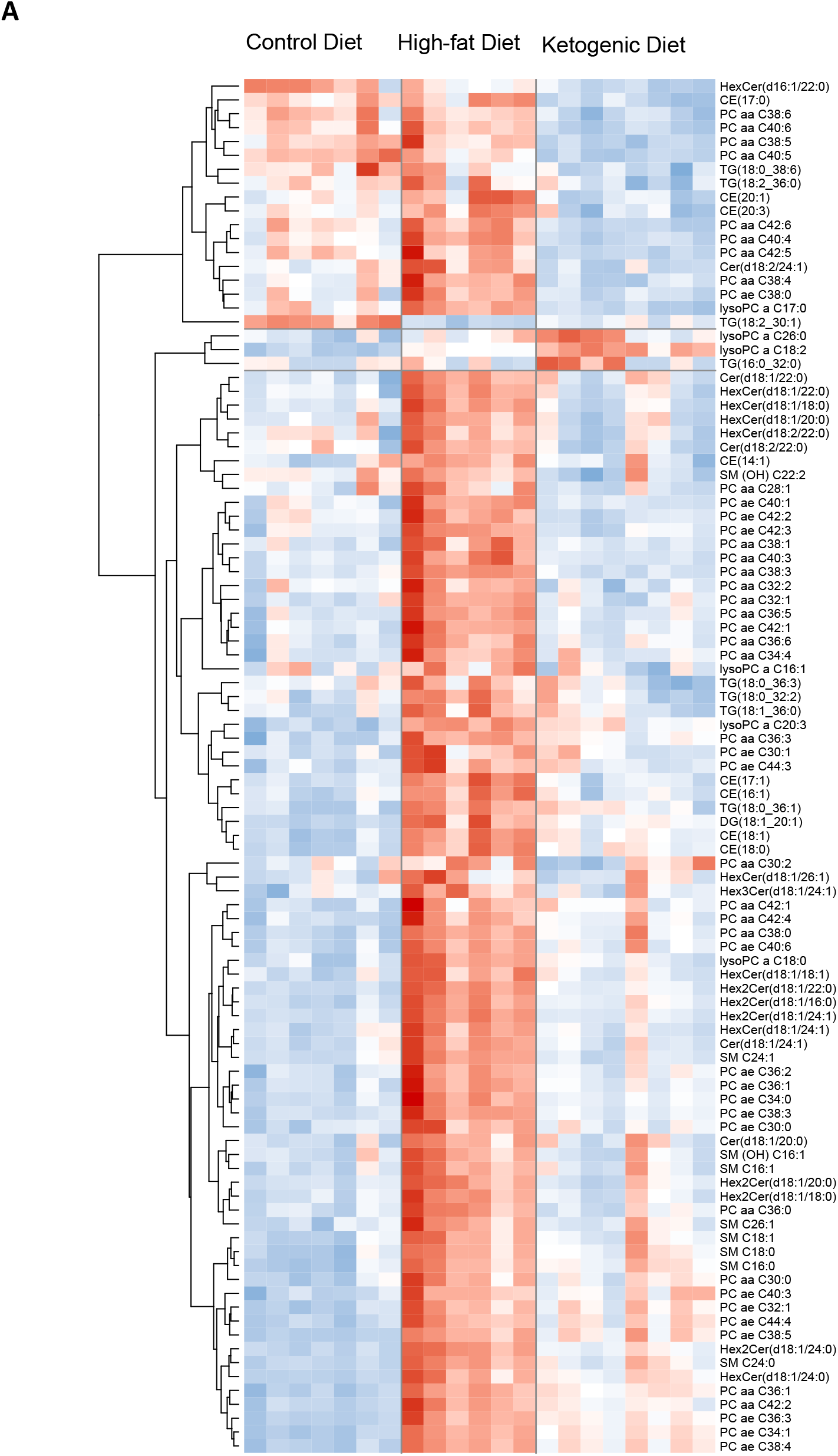
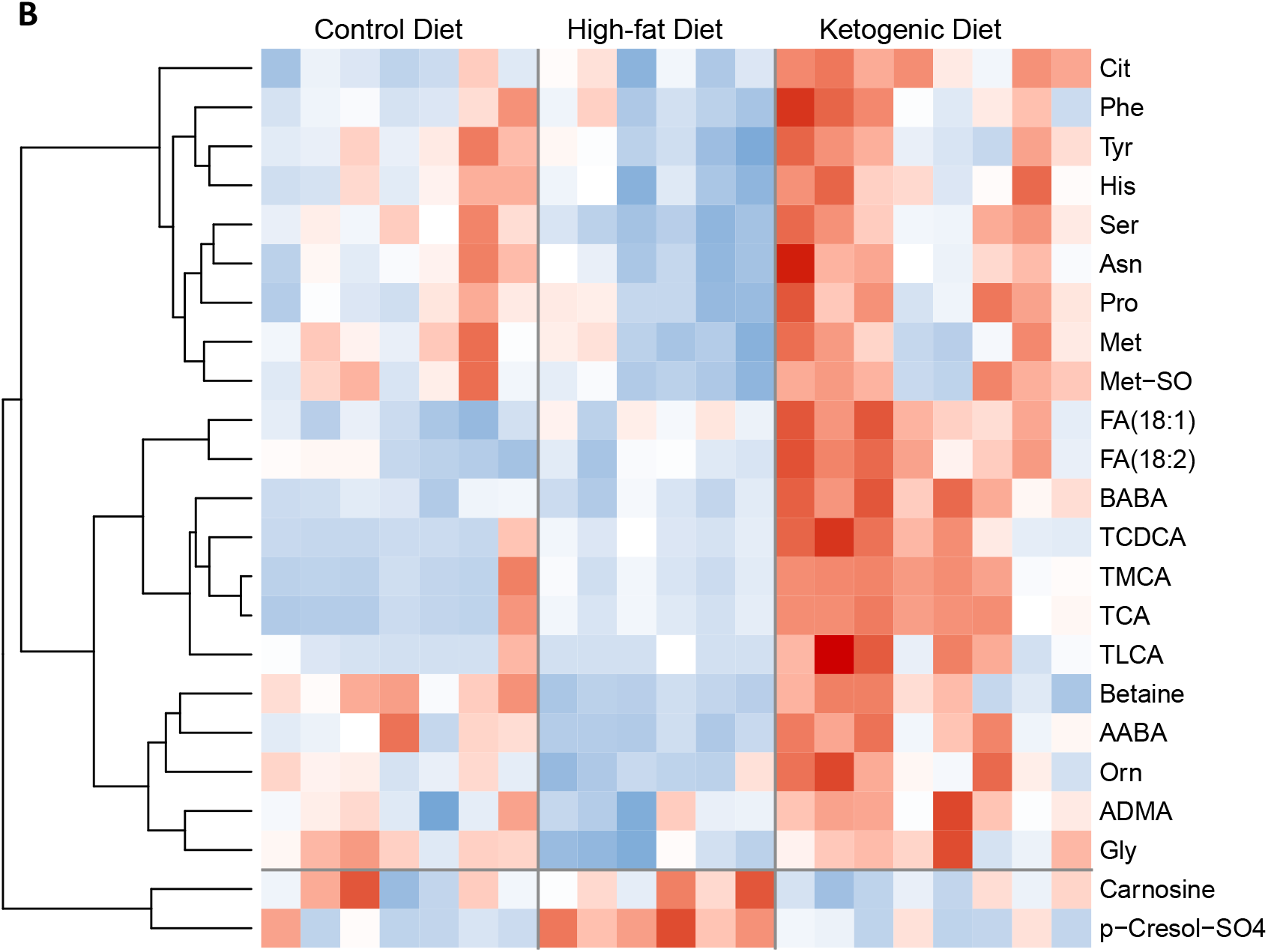
Heat maps showing all 99 complex lipids (A) and all 23 small molecules (B) significantly (FDR <0.05) altered by the diets.

**Figure 5.**
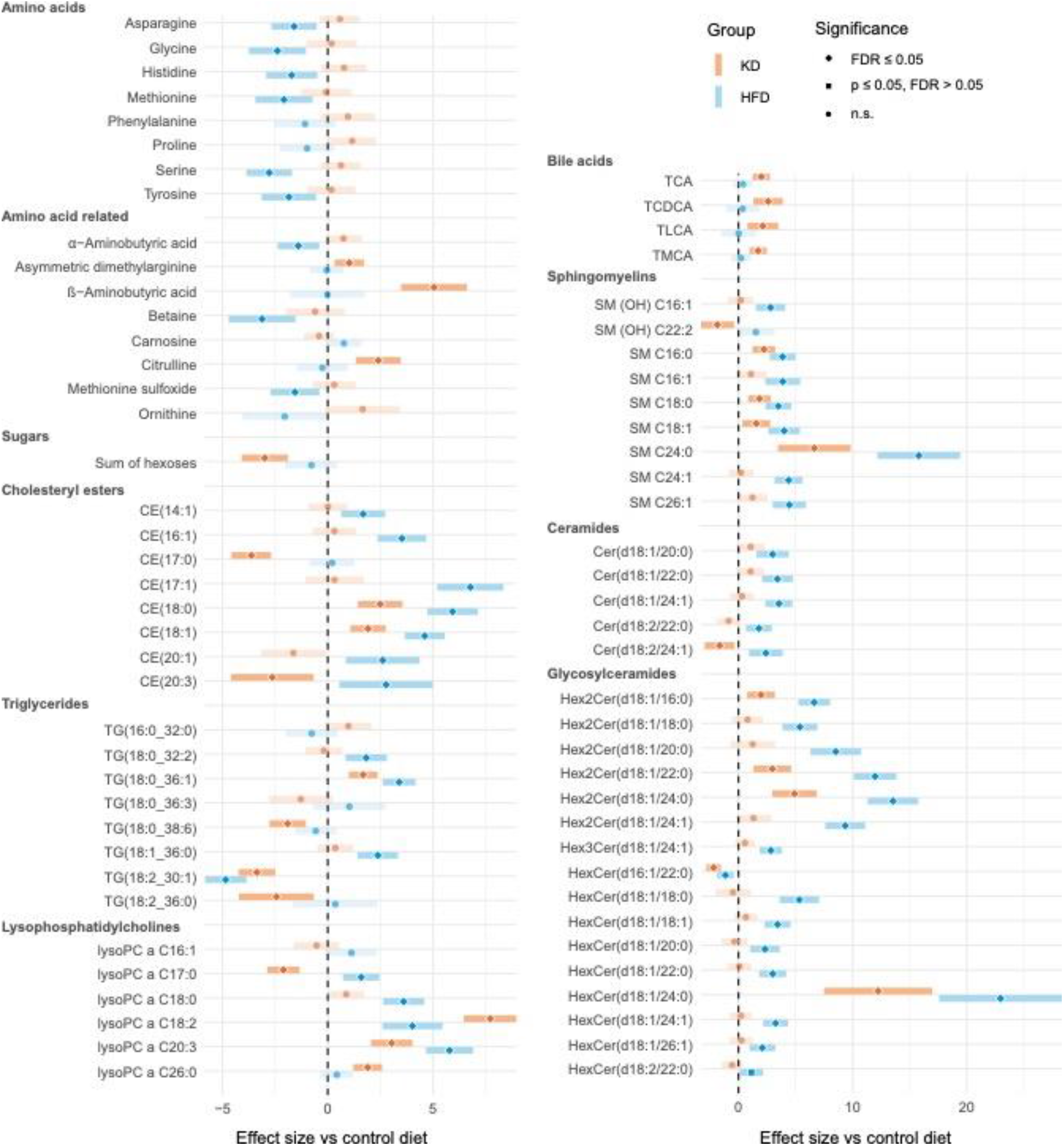
Forest plots of metabolites represented as normalized regression coefficients with 95% confidence intervals. The black line represents a zero effect of the control diet with significant differences (FDR <0.05 or p-value <0.05) indicated for either HFD or KD. Regression models include covariates as detailed in the Methods.

We also examined the differences in 192 metabolic indicators determined in all groups of mice. These indicators are sums or sums and ratios of metabolites from the Quant 500 analysis and cited as biomarkers that can reveal information on specific pathway activity and metabolic flux. We found 74 metabolic indicators that were significantly different (FDR <0.05) in the KD compared to the HFD mice. These indicators showed a similar pattern as observed by the individual metabolites, with the KD versus HFD mice showing an accumulation of several indicators related to the metabolism of the amino acids but a decrease in the sum or ratio of several lipid species as well as indicators of gut microbial activity (Figure 6; Supplementary Table S2).

**Figure 6.**
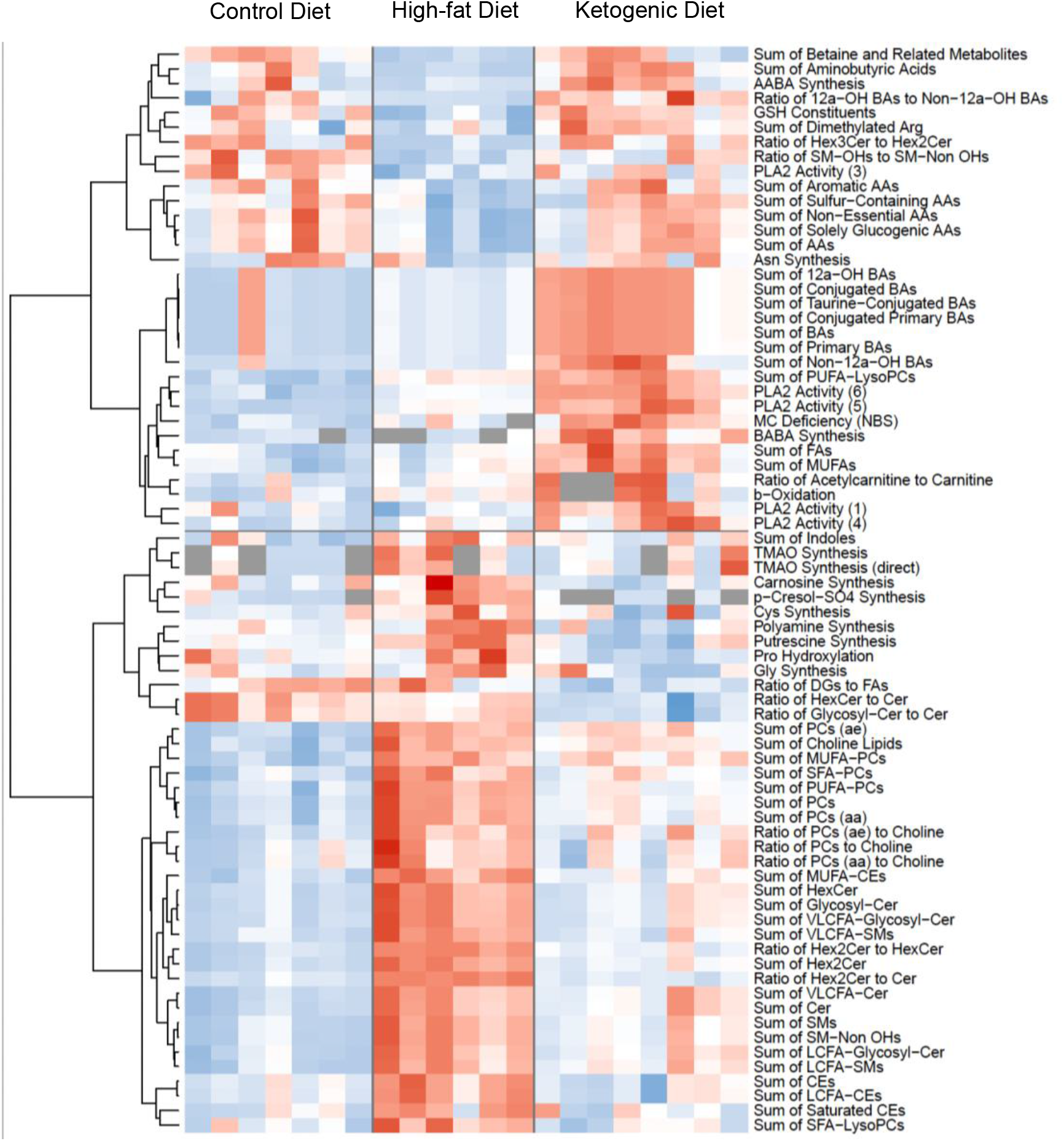
Heat map of the 74 metabolic indicators significantly (FDR <0.05) altered by the experimental diets. Increasing intensity of red color indicates higher concentrations of metabolites and increasing intensity of blue color indicates decreased concentrations of metabolites.

Lastly, a cluster analysis was performed in the 123 significantly altered plasma metabolites between KD and HFD-fed mice. The results were used to map out the identified metabolites into specific chemical clusters. The results confirmed that the KD significantly decreased several lipids species while increasing the concentration of hydrophilic metabolites (Figure 7; Supplementary Table S3).

**Figure 7.**
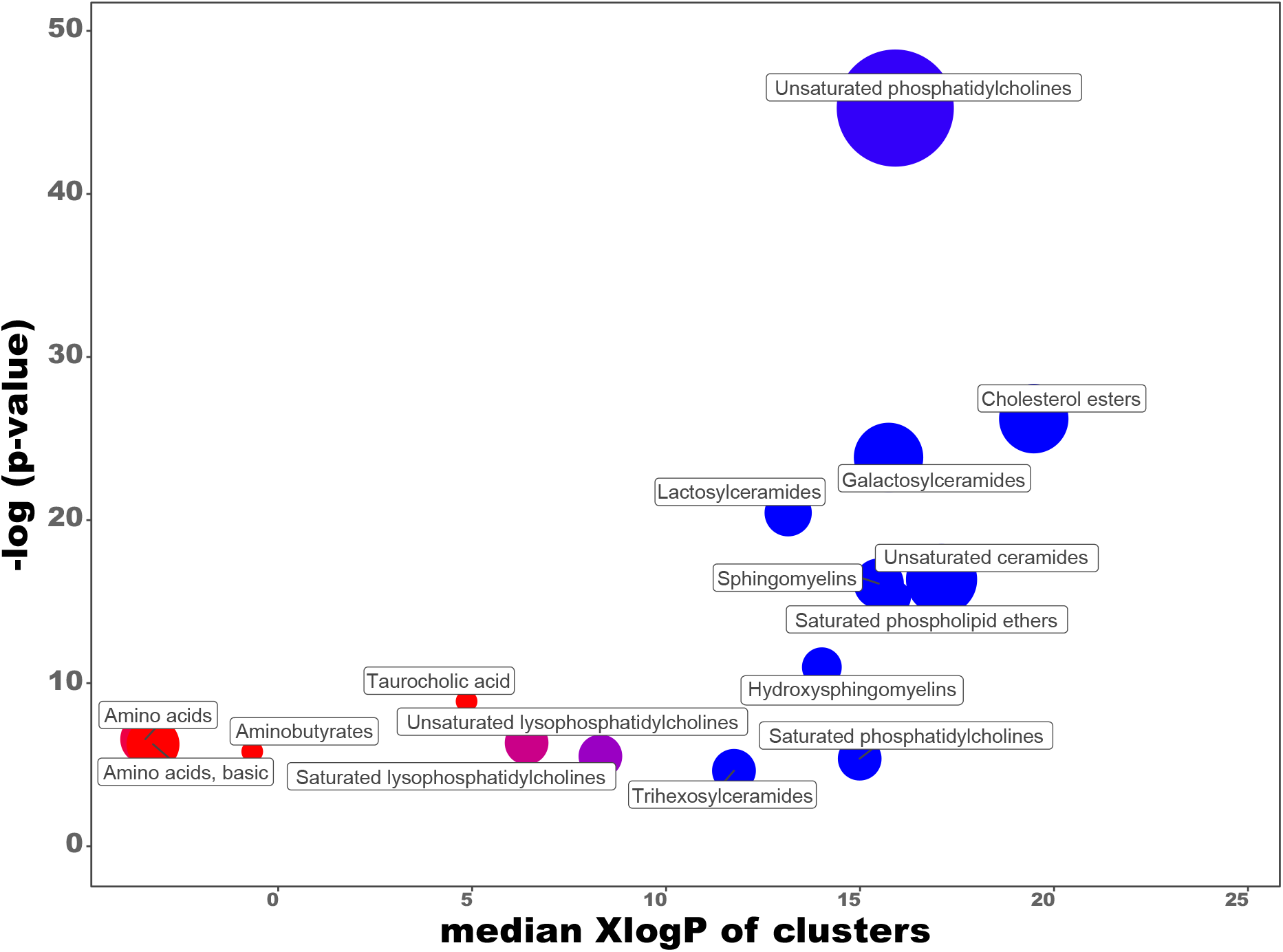
Chemical clusters of metabolites (C) altered by the ketogenic diet, when compared to the high fat diet. Increasing intensity of red color indicates higher concentrations of metabolites and increasing intensity of blue color indicates decreased concentrations of metabolites.

## Discussion

In recent years, KDs have been considered a promising strategy for weight management in humans ^8,29,30^ and carbohydrate restriction may be more effective than fat restriction for obesity treatment ^31^. Previous studies in mice have also reported the same anti-obesogenic effect for KDs, despite the high high-fat content ^32-35^. In this study, although caloric intake was similar in mice-fed diets with a high-fat content, i.e., KD and HFD, KD-fed *ApoE*^—/—^ mice weighed significantly less than HFD mice.

KDs stimulate endogenous ketogenesis, thus yielding high levels of BHB ^13^. The plasma concentration of the major ketone body, BHB, was significantly elevated in KD-fed mice compared to the other two mice groups and within the range reported by others ^13,34^. This finding confirms the presence of nutritional ketosis in KD-fed mice, and it indicates the reliability of this dietary mouse model in understanding the effect of nutritional ketosis on vascular homeostasis.

Atherosclerosis is a disease characterized by low-grade, chronic arterial wall inflammation ^36^. Notably, the efficacy of anti-inflammatory therapy in human atherosclerosis has been recently validated ^37,38^. Several pro-inflammatory cytokines are atherogenic in humans and mice ^39-41^. These include MCP-1, implicated in infiltrating immune cells into the vessel wall, an essential component of atherosclerosis progression ^42^. Another example is IL-6, which plays a central role in propagating the downstream inflammatory response that sustains atherosclerosis progression ^43^. Lastly, TNF-α is a crucial cytokine that facilitates the influx of inflammatory cells into the vessel wall, thus promoting the advancement progression of vascular lesions ^41^.

Therefore, we investigated the effect of the experimental diets on the circulating levels of MCP-1, IL-6, and TNF-α. The results showed that the HF diet promoted a significant systemic elevation of all three cytokines compared to the control diet. This observation agrees with the well-established effect of dietary fat on systemic inflammation ^27,44^. As previously mentioned, the KD contained approximately 80% fat, whereas the fat content in HFD was half (40%).

Interestingly, despite the KD’s higher fat content compared to the HFD, plasma from KD-fed animals presented significantly lower levels of the investigated cytokines than the HFD, thus suggesting that nutritional ketosis was able to attenuate the fat-induced inflammatory effect. In accordance with the diminished systemic inflammation detected in mice fed a KD, the results showed that KD mice presented significantly less aortic atherosclerosis burden than HFD mice. Furthermore, the atherosclerotic lesions were localized in the same aortic regions, the aortic arch and brachiocephalic artery, in all three groups. So, the differences between HFD and KD aortas in these highly susceptible regions were even more pronounced. Previous studies, including ours, in which HFD with very low-carbohydrate contents were fed to murine models of atherosclerosis reported the atherogenic effects of these diets ^45-47^. However, in these studies, the vascular toxicity of the diets was compared with a control diet, thus more in line with the AMDR in humans. In the present study, we also observed that the KD’s volume of vascular lesions was increased compared to the controls. However, when comparing the aortic plaque burden with the one associated with the HFD, which better represents the average American diet, the extent of atherosclerotic lesions was significantly lower and thus not parallel to the amount of dietary fat.

This intriguing observation suggests that the presence of nutritional ketosis alleviated fat-induced vascular toxicity. Increasing evidence suggests a positive impact of nutritional ketosis on vascular physiology and homeostasis in humans ^48-50^. For example, a recent experimental human study showed that carbohydrate restriction benefits multiple components of the metabolic syndrome, a major CVD risk factor ^51^.

Preclinical experiments on animal models are essential to understanding CVD progression mechanisms. *ApoE*^—/—^ mice have been widely used as an animal model to study the pathophysiology of atherosclerosis due to their striking similarities with humans in the molecular mechanisms that lead to endothelial dysfunction and vascular plaque formation ^14,52,53^. However, the translational value of an animal model strongly depends on its ability to robustly reproduce important functional, structural, and molecular pathological features of human disease. *ApoE*^—/—^ mice are one of the available animal models of human atherosclerosis that better mimics the human metabolite signature ^54^. Thus, to better understand the biological processes related to the influence of nutritional ketosis in the development and progression of atherosclerosis, we assessed the metabolomic signature in the plasma from mice in the different experimental groups. As mentioned, the fat content in the HFD is 40%, whereas the KD is 80%. However, significantly lower levels of several lipid species in various lipid classes were observed in the KD group versus HFD. The most significantly affected pathways by atherosclerosis progression in *ApoE*^—/—^ mice are glycerophospholipid and sphingolipid metabolisms ^55^. In accordance, here we observed an accumulation of several phosphatidylcholines, ceramides, hexosylceramides, and sphingomyelins in the group of HFD-fed mice, which showed the highest plaque burden.

Notably, however, a significantly decreased concentration of these lipid metabolites was observed in the KD group, suggesting that this favorable metabolic profile resulted from using fat as the primary energy source. A similar pattern was indicated by the decreased metabolic indicators and sums/ratios of several lipid species in the KD-mice compared to the HF group. Interestingly, however, the sum of fatty acids (FAs), including monounsaturated fatty acids (MUFAs), were found to be significantly accumulated in the animals under nutritional ketosis, i.e., under KD, probably reflecting its dietary origin and high concentration in the diet.

Ceramides, a class of sphingolipids, are constituents of every cell membrane and present with a wide range of cellular actions, including at the vascular level ^56^. For example, endothelial dysfunction is the first step in the establishment of atherosclerosis and increased ceramides have been implicated in the disturbance of endothelial homeostasis by different mechanisms including elevated oxidative stress. In addition, ceramides species have been shown to initiate apoptosis and can inhibit antiapoptotic mediators ^57^ and cell differentiation ^58^. Furthermore, increased ceramides have been implicated in inflammation and vasoconstriction, and can activate the NLRP3 inflammasome to induce apoptosis in macrophages and adipose tissue ^59^. Accordingly, over the past few years several human studies reported the association of circulating ceramide levels with adverse cardiovascular events ^56^. In *ApoE*^—/—^ mice, an increase in C18-24 ceramide levels favored atherogenesis, while C16:0 ceramide has an opposite function ^55^. In agreement with previous studies, several C18 ceramides species were significantly increased in the HFD mice, when compared to the KD group, which presented less plaque burden than the other group of mice in this study.

The presence of BHB is a consequence of enhanced hepatic fatty acid β-oxidation that occurs as a response to the very low carbohydrate levels in the KD. Carnitine transports activated long-chain FAs, in the form of acylcarnitine, from the cytosol into the mitochondrion and is, therefore, essential for FA β-oxidation. Consistent with the shift in energy metabolism toward fatty acid oxidation caused by the KD, the ratios of acyl-carnitines, including the short-chain species, to free carnitine was significantly higher in KD than in HFD mice ^60^.

The levels of several non-lipid metabolites were also affected by the experimental diets.

For example, despite the same source and amount of protein in the KD and HFD, mice under nutritional ketosis accumulated several plasma amino acids suggesting a KD-induced alteration of the protein metabolism that resulted in significantly higher levels of total amino acids, as well as aromatic, sulfur-containing, nonessential and solely glucogenic amino acids. The constituents of the major antioxidant glutathione, i.e., the amino acids glutamate, glycine, and cysteine, were also significantly increased by the KD, suggesting a more favorable redox status in the animals under nutritional ketosis. KD-fed mice developed less extensive atherosclerosis than HF mice, which may have been contributed to, in part, by a decreased oxidative stress in the KD mice. The pathologic process of atherosclerosis begins with the trapping of oxidized lipids in the vessel wall. Thus, oxidative stress is a significant contributor to the establishment and progression of the atherosclerotic process ^2,61^. Ketosis is an evolutionarily ancient metabolic pathway that might confer benefits by several mechanisms, including modulation of oxidative damage and cellular redox state ^62^.

The effect of the experimental diets used in this study on the systemic methylation index, the plasma adenosylmethionine to adenosylhomocysteine ratio, has been previously evaluated in *ApoE*^—/—^ mice ^16,17^. A significantly decreased plasma AdoMet:AdoHcy ratio was detected in KD-fed mice but not in HF-fed mice, versus controls, thus suggesting the existence of systemic hypomethylation under nutritional ketosis. In accordance with this possibility, we observed an accumulation of betaine in the KD-fed mice versus the HFD mice, suggesting the underutilization of this methyl donor in the folate-independent remethylation of homocysteine to methionine ^63^. Moreover, the levels of more than forty phosphocholines were significantly decreased in the KD-mice compared to the HFD mice. Studies have shown that, in fact, approximately 30% of PC is synthesized in the liver by 3 transmethylation reactions catalyzed by phosphatidylethanolamine N-methyltransferase (PEMT) ^64^, which activity can be impaired by a decreased AdoMet:AdoHcy ratio. Noteworthy, the lack of PEMT protected against diet-induced atherosclerosis in two mouse models ^64,65^.

Bile acids (BA) are signaling molecules that regulate lipid and energy metabolism ^66^. The two primary BA receptors, the membrane TGR5, and the nuclear FXR are implicated in cardiovascular disease pathogenesis ^67^. Accordingly, rodent models of atheroma documented a powerful anti-atherosclerotic effect of BA ^67^. BA are synthesized from cholesterol in the liver and stored in the gallbladder to be secreted in the duodenal lumen upon fat intake to facilitate digestion ^68^. Primary BAs are also conjugated in the liver with the amino acid glycine or taurine to enhance water solubility. When the BAs reach the colon, the microbiota transforms primary BAs into secondary BAs. In this study, compared to the group fed the HFD, the plasma concentration of several individual, primary, and conjugated BAs was increased in the KD-fed group, which suggests (that more BA in KD group might helped in fat elimination from the body) the role of BAs in terms of fat elimination.

Increased plasma levels of asymmetric dimethylated arginine (ADMA) have been associated with atherosclerosis and negative cardiovascular events in humans and mice ^36,69^. ADMA is an endogenous inhibitor of nitric oxide synthase (NOS), the enzyme responsible for nitric oxide, a major regulator of vascular homeostasis. However, in our study, ADMA levels were higher but plaque burden was lower in the mice fed KD versus the HFD group, an observation that does not agree with the previously reported atherogenic effect of ADMA.

Importantly, some experimental clinical studies failed to prove the causal role of increased ADMA concentrations in inflammation, a major player in the development of atherosclerosis ^69^.

Lastly, three metabolic indicators suggested differential microbial activity induced by the experimental diets. The ratios of trimethylamine N-oxide (TMAO) to the sum of betaine, free carnitine, and choline, as well as to choline are indicators of TMAO synthesis; TMAO is a metabolite associated with CVD and its precursor, trimethyl amine, TMA, is produced by the gut bacteria ^70,71^. The ratio of p-cresol sulfate to tyrosine indicates tyrosine degradation by gut bacteria activity ^72^. Compared to HF-fed mice, KD-fed mice presented decreased values for these three metabolic indicators, an observation that warrants further investigation. Notably, recently published works reported that KD-associated microbiome-metabolomics changes resulted in an anti-inflammatory activity ^73,74^.

In conclusion, nutritional ketosis induced by the KD versus HFD was associated with anti-obesogenic and anti-inflammatory systemic effects, significant metabolic changes that affected the levels of several lipid species, amino acids, and bile acids, and promoted an atheroprotective phenotype. It remains to be determined if our findings in mice on the metabolic and vascular effects of KD and HFD translate to the same extent in humans. This study provides the rationale to support clinical trials to understand further the underlying mechanisms and the anti-atherogenic effect of KD’s in humans.

## Funding

This work was supported by the following funding sources at Pennsylvania State University: Department of Nutritional Sciences, the High-Field Magnetic Resonance Imaging Facility, and the Huck Institutes of the Life Sciences.

### Institutional Review Board Statement

The study was conducted according to the guidelines of the Declaration of Helsinki and approved by the Institutional Animal Care and Use Committee of Pennsylvania State University, which specifically approved this study (PRAMS#201747911 to R.C.) in November 2018.

## Acknowledgments

The authors wish to thank Courtney A. Whalen and Floyd J. Mattie (Penn State University, University Park, PA, USA) for expert technical assistance and Isabel Tavares de Almeida (University of Lisbon, Lisbon, Portugal) for her valuable support.

## Supplementary Tables

S1. List of metabolites included in the MxP Quant 500 assay. Statistics for control diet (CD) and ketogenic diet (KD) in comparison to high fat diet show p-values and FDR (≤0.05 indicated in yellow) from regression analysis and log2 fold change of group means. NA = values not available due to exclusion of metabolites during filtering.

S2. List of metabolic indicators included in the MxP Quant 500 assay. The indicators were calculated by MetIDQ software using Metabo*INDICATOR* formulas. Statistics for control diet (CD) and ketogenic diet (KD) in comparison to high fat diet show p-values and FDR (≤0.05 indicated in yellow) from regression analysis and log2 fold change of group means. NA = values not available due to exclusion during filtering.

S3. ChemRICH analysis showing identified clusters, assignment of metabolites into the clusters, and cluster statistics including p-values and FDR (≤0.05 indicated in yellow).

